# Vive la difference: why, how, and what trait combinations improve functional community ecology

**DOI:** 10.1101/2023.09.29.560181

**Authors:** Alexandra Engler, Dylan J. Fraser, Pedro Peres-Neto

## Abstract

Analyses of functional trait variation across ecological communities offer valuable insights into the factors that influence and predict trait composition, deepening our understanding of community structure. These analyses also shed light on patterns of trait dispersion, including underdispersion and overdispersion, which are often linked to the mechanisms underlying community assembly. However, striking a balance between characterizing complex phenotypes through the integration of comprehensive set of traits while ensuring the preservation of the distinct signals of overdispersion and underdispersion can pose significant challenges. This is because some trait combinations can result in high functional dispersion or overdispersion, while others may lead to underdispersion within the same community. In this study, we provide a rationale for the importance of trait selection in community ecology and develop a framework that optimizes patterns of functional trait variation among communities. We explored the responses of six trait sets along large-scale environmental gradients, using ∼700 lake-fish communities in Ontario (Canada). We started by adopting the conventional approaches of combining all traits as well as leveraging prior knowledge in fish biology to group traits that are functionally related (Diet, Morphology, Temperature Preference). We contrasted these approaches with a novel computational method to select combinations of traits that maximise overdispersion and minimise underdispersion. We found that separating traits according to their functions or by their patterns revealed signals in communities that appeared random when all the traits were pooled together. Combining our computational approach with environmental models revealed that functional patterns could be very well explained by multiple environmental predictors (model R2 up to 0.7). Our study also outlines future selection procedures to further uncover more complex patterns of species associations based on their traits.

## INTRODUCTION

Functional ecology provides a robust foundation for understanding species coexistence and ecological community assembly (Götzenberger et al. 2012; Münkemüller et al. 2012; Adler et al. 2013; Munoz et al. 2023). Functional traits describe the ecological requirements of individuals or species and approximate various components of their niches. Functional metrics, such as functional dispersion, provide valuable insights into assembly mechanisms (Münkemüller et al. 2012; Adler et al. 2013), enabling us to better understand the factors that influence and predict trait composition within ecological communities. Environmental selection reduces functional dispersion, favoring species with specific trait sets for survival and persistence (Mouillot et al. 2007; Adler et al. 2013; Mason et al. 2013). Conversely, competition leads to high functional dispersion as similar species exclude each other (Adler et al. 2013; Mason et al. 2013). However, interpreting functional dispersion patterns alone may not always provide clear inferences (Cadotte & Tucker 2017). For example, competition may lead to local communities composed of species that share traits (clustering), resulting in the wrong inference of environmental selection (Mayfield & Levine 2010).

Large-scale observational studies are critical for understanding the sources of trait variation in ecological communities, despite the challenges related to mechanistic inference. These studies increase our predictive abilities to predict patterns that can be contrasted and validated across multiple systems and spatial scales (McGill et al. 2006). Unlike species composition, functional traits are often shared across multiple global ecosystems, thus serving as common predictive “currencies” across ecosystems. Additionally, the strong association between environmental features and trait dispersion provide valuable insights on how environments influence local demographic variations among species (Cadotte & Tucker 2017). Finally, community trait values change as a function of spatial and environmental extents, offering a solid basis for explaining common trait-environmental trade-offs observed in nature that are difficult to explore through experiments (Kneitel & Chase 2004).

While inferring trait dispersion patterns, it is common to consider mutually exclusive mechanisms such as environmental selection and species interactions. However, it is important to note that multiple mechanisms can act within single local communities (Mouillot et al. 2007; Mason et al. 2013; Trisos et al. 2014). This acknowledgement emphasizes two key points. First, different mechanisms may select different sets of traits within local communities (e.g., some trait combinations can result in high functional dispersion or overdispersion, while others may lead to underdispersion within the same community). Second, combining different sets of traits weakens our ability to detect distinct community trait patterns (Cornwell & Ackerly 2009; Côte et al. 2019), particularly in highly heterogenous environments (e.g., lakes, streams, ecotones). This is a critical consideration because most studies combine multiple traits when estimating single metrics of functional dispersion. Selecting and aggregating traits to study assembly mechanisms is a complex task. Using a single trait may not capture the complexity of species’ phenotypes, while an excessive number of traits can introduce redundant information (Zhu et al. 2017; Mouillot et al. 2021).

We analyse a comprehensive dataset containing hundreds of lake-fish communities across a large environmental gradient to investigate how and what different methods of aggregating trait sets improve the detection and prediction of community functional patterns. Lake-fish communities, like islands, impose net boundaries that limit species to dispersal (Magnuson et al. 1998) compared to continuous terrestrial ecosystems, streams or marine areas. As a result, local fish populations experience strong environmental or biotic selection (competition and predation) due to restricted movement among lakes (Olden et al. 2001). In fish communities, diet traits (e.g., jaw gape, trophic level, gut length) are expected to exhibit greater dispersion (overdispersion) when resources are limited due to competition. This is expected particularly in lakes in which environmental filtering is weak and that can support large and diverse communities. Conversely, traits related to species movement (e.g., fin positions and sizes) are more likely to undergo stronger environmental selection (Villéger et al. 2017; Côte et al. 2019) and show underdispersion when unfavorable abiotic conditions restrict the presence of numerous species. These lines of argumentation reinforce the notion that different combinations should exhibit distinct responses, and associated patterns to environmental variation, as well as highlight potential synergies and trade-offs among trait sets (Cornwell & Ackerly 2009; Lopez et al. 2016).

## METHODS

### Data and sampling design

The Ministry of Northern Development, Mines, Natural Resources and Forestry have sampled 707 lakes across Ontario for their Broadscale Monitoring Program (OBMP, Lester et al. 2020). Lakes were sampled in summers (June to September) from 2008 to 2012 along a large latitudinal and longitudinal gradient (Figure 1). The OBMP uses stratified sampling: lakes were chosen randomly among lakes of similar sizes within the same region. They spread over three primary watersheds and 21 secondary watersheds.

**Figure 1.**
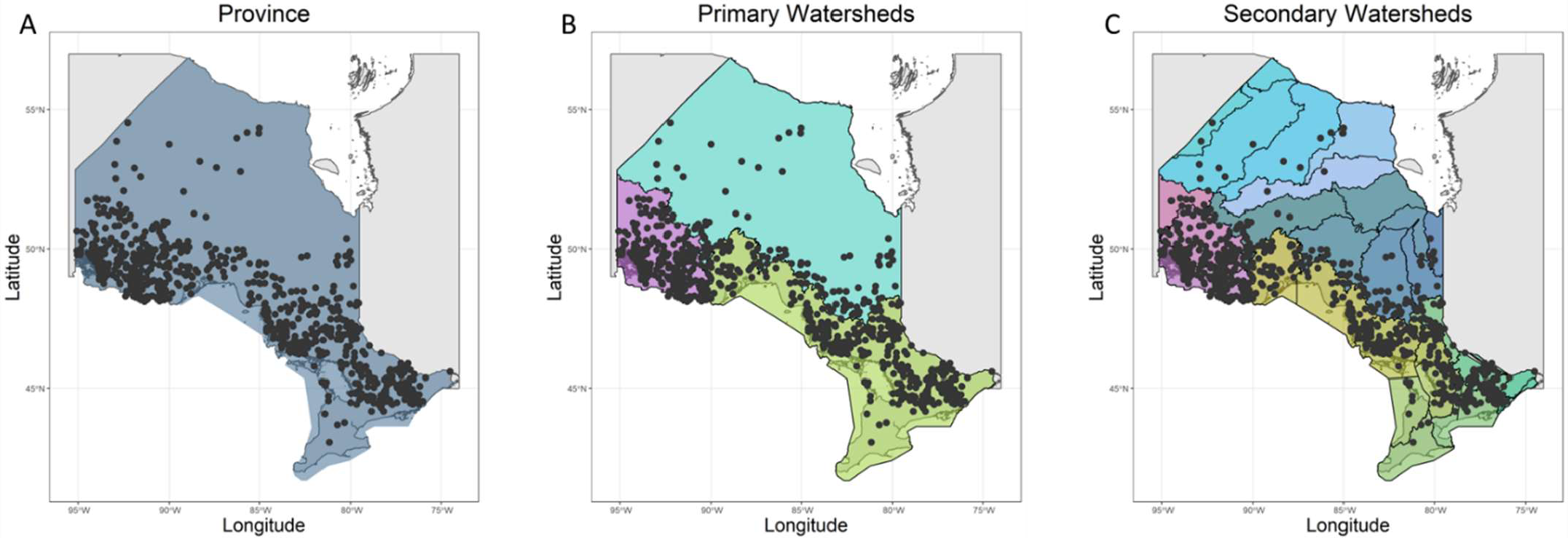
Maps of the spatial definition of the different species pools for lake-fish communities of Ontario, Canada. The polygons represent the geographical boundaries with which the species pools were defined. The points are the sampled lakes.

Each lake was sampled using a standardized design. Fish were caught at different depths with two multi-mesh gill nets; one small mesh net, which stretched from 13 to 38 mm, and one large mesh net that stretched from 38 to 107 mm. The number of nets per lake was set based on lake size and average depth. Nets were set overnight, following the standards for freshwater community sampling (see Appleberg 2000 for general fish sampling; and Arranz et al. 2022 for further details on the OBMP sampling framework). For each lake, every fish was identified at the species level. The number of species ranged from 2 to 25 per lake. Only lake communities with more than 3 species were considered for analysis (696 out of 707) (Table 1).

**Table 1.** Distribution of lake community (replication) across spatial scales used to infer species pools.

| Scale of inference | Scale at which the factor of interest was applied | Number of observations at the appropriate scale |
| --- | --- | --- |
| Community (lake) | Province | 696 lakes and 1 Province |
| Community (lake) | Primary Watersheds | 696 lakes and 3 Primary Watersheds |
| Community (lake) | Secondary Watersheds | 696 lakes and 21 Secondary Watersheds |

Lake environment was characterized by 86 variables recorded for each lake, which we then divided into six environmental categories (details in Appendix 1: Table S2): Climate (e.g. minimum monthly air temperature (MnMonTP), cumulative temperature below or above 0°C), Water quality (e.g., Nitrate or Potassium concentration), Hydro-morphology (lake area, littoral perimeter), Watershed and spatial characteristics (e.g. altitude, lake age, Tertiary Watershed elevation), and Fishing Activities (e.g., angling pressure, number of boats during summer). Before linking trait dispersion patterns to environmental variation via regression, we normalized the variables with the function BestNormalize (package BestNormalize, Peterson 2021) and standardized (mean=0, variance=1) them. As some variables were colinear, we summarized their variation based on axes from a principal component analysis (PCA) for each environmental category. The number of axes was set to keep no less than 80% of the variance from each environmental category.

### Species traits and selection of trait sets

We selected 15 traits (Table 2, details in Appendix 1: Table S1) representing different aspects of species ecology and grouped them into four categories: Diet, Morphology, Temperature preference, and a set with all traits together (Table 2). The traits data were extracted from FishBase (Froese & Pauly 2019). As in most trait-based studies, these traits are summarized at the species level without considering potential intraspecific variation. Prior to analyses, traits were standardized to mean zero and unity variance to avoid artificially putting more weight on one trait over others.

**Table 2.** List of fish traits by function based on Villéger et al. (2017) and Schleuter et al. (2012) and the expected patterns.

| Function | Traits | Expectations |
| --- | --- | --- |
| All traits pooled together | 15 numerical traits | Random patterns |
| Diet | Length, Eye diameter, Trophic level, Preorbital length | Mainly over-dispersion: competitive interactions |
| Morphology | Aspect ratio, Standard length, Fork length, Preanal length, Predorsal length, Prepelvic length, Body depth, Head length | Mainly under-dispersion: environmental filtering |
| Temperature tolerance | Minimum temperature, maximum temperature, interval between minimum and maximum temperature | Mainly under-dispersion: environmental filtering |

Here we explored two approaches to determine how trait combinations improve (or decrease) our ability to detect and predict functional variation among communities. First, we used *a priori* categories based on fish biology, representing functional traits that enable species to perform specific functions. Following Villéger et al. (2017), traits were grouped into three categories: diet, morphology, and temperature preference (Table 2). To ensure equal influence among categories when combining all trait sets, we used Gower’s (1971) standardization for each category separately. This standardization, which, to our knowledge, has not been used in trait studies, equalizes the total sum of squares of each set to one (see Peres-Neto & Jackson, 2001). The second approach involved a novel exploratory framework that generates trait sets optimized for under or overdispersion patterns based on the following forward selection approach: 1) Select the trait that, on average across lakes, had the smallest standardized community dispersion metric (i.e., the most underdispersed trait; (see metric and null models below), T1; 2) Pair each remaining trait with T1 and select the next trait that further reduced the community dispersion metric the most, T12. 3) Repeat step 2 (i.e., T123, T1234, and so on); 4) When no further reduction is possible, the algorithm stops, and we keep the selected traits from the previous step. Repeat the same procedure considering overdispersion instead. Although our selection procedure aims at increasing pattern detection in an exploratory way, it should also allow uncovering complex ways in which different functions are associated in the assembly process. To this end, we 1) conducted network analysis based on the correlation of the community dispersion metric values (see below) across lakes; and 2) calculated the overall strength of these correlations as the average absolute correlation values among traits. These analyses allow one to assess how traits are integrated into the assembly process. The absence of strong positive associations in community functional trait metrics suggests differential roles of traits in species sorting and environmental filtering. Conversely, negative correlations between community functional trait metrics (e.g., overdispersed trait negatively correlated with an underdispersed trait) imply the simultaneous influence and interaction of environmental filtering and niche differentiation in the assembly process. Finally, the correlation analysis enabled us to determine unexpected ways in which traits that are assumed to differ dramatically in their functions (e.g., temperature versus morphological traits) can be correlated (positively or negatively) in the way they contribute to community assembly. Correlation networks of community functional trait metrics are an unexplored venue with the potential to improve further our understanding of the processes underlying community assembly.

## Statistical analyses

### Quantifying multivariate trait patterns within local communities

The starting point of community-trait analyses is to calculate species pairwise distance among species based on individual or combination (sets) of traits as outlined earlier. We used the Gower distance (Gower 1971) given its ability to handle missing trait values for a few species. The resultant pairwise matrix was then used to estimate functional dispersion for each community (lake). We calculated functional dispersion (diversity) based on Hill numbers (Chiu & Chao 2014), as it is one of the most robust in detect under- and/or overdispersion (Mammola et al. 2021):

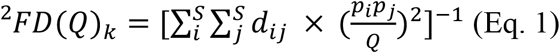

where 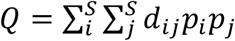 (i.e., Rao’s quadratic entropy; sum of the weighted pairwise distance in the community), *S* is the total number of species, *d*_ij_ is the pairwise (Gower) distance between species *i* and *j, p*_i_ and *p*_j_ are weights for species *i* and *j*, respectively (here p is either 1 or 0 if the species is present or absence in a particularly lake, respectively) and k is an index to indicate the *k*^th^ lake (i.e., each lake has its own dispersion metric).

As Rao’s is a metric of absolute functional dispersion of the communities, *FD*_*Q*_ increases as a function of the number of species in local communities (lakes) and, as such, lakes cannot be compared directly. *FD*_*Q*_ could be higher for a lake comprised of many randomly selected species than when compared with a lake with a smaller number of species that are not randomly selected. To make *FD*_*Q*_ comparable across local communities, we used its customary Standardized Effect Size (SES) based on Hill number:

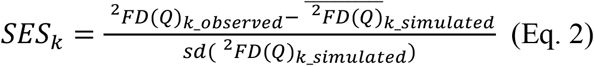

where *SES*_*k*_ is the standardized effect size for the *k*^th^ lake and ^2^*FD*(*Q*)_*k_observed*_ is the observed functional dispersion for that lake, 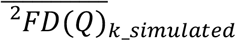 and *sd*(^2^*FD*(*Q*)_*k_simulated*_) are the average and the standard deviation, respectively, expected for randomly assembled communities (simulated under a particular null model; see below). Negative *SES* in lakes imply species have a high similarity than expected by chance (underdispersed), while positive *SES* indicate that their species are more distinct than expected by chance (overdispersion). We also estimated the p-value of each SES value following Eq.3:

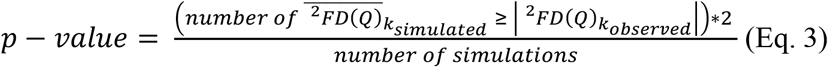

### Null models for generating randomly assembled communities

SESs were computed by creating null communities through the random sampling of the same number of species as present in each lake from a species pool assumed to represent potential colonizers. We generated 999 randomly assembled communities for each lake. Recognizing that the choice of species pools in null model analyses can determine the spatial scales at which communities are structured (Lessard et al. 2012), we considered pools across three different spatial scales: 1) Province scale, encompassing all species across all lakes in the dataset, 2) Primary watersheds, and 3) Secondary watersheds, in which species pools were defined separately for each watershed level (Figure 1).

### Environmental analysis of variation in functional trait patterns

In Ontario, the strongest environmental gradient is known to follow a south-north trajectory (e.g., Henriques-Silva et al. 2019). We started by fitting a Generalized Additive Model (GAM) to detect the strength of non-linear latitudinal trends in SESs for each spatial scale (species pools). We fitted a GAM model for each trait set and spatial scale using latitude as a predictor. The maximum number of splines (k) were set at 20 and model regularization was used to estimate parameter that best fitted the SES based on different spatial scales and set of traits. The smoothing parameters were estimated with the REML (restricted maximum likelihood) method.

The next step was to understand how SES varies as a function of environmental gradients. To do so, we fitted a GAM model for the different SES (spatial scale and trait combinations) against the PCA for axes (see section *Data and sampling design*) for each environmental category. The number of splines for each estimator was set in the same way as described above. We then extracted the adjusted R^2^ to assess how different set of predictors were able to explain patterns of community trait variation (SES).

All analyses were performed in R (R Core Team 2020). The Gower’s distance and trait metrics were calculated with the package ‘FD’ (Laliberté et al. 2014) and ‘mFD’ (Magneville et al. 2022). Null models and trait combinations that maximized (or minimized) SES were computed via a custom-made function. GAMs were fit with the package ‘mgcv’ (Wood et al. 2016).

## RESULTS

### Identifying combinations of traits and correlations among functional dispersion patterns

Traits were combined either *a priori* (i.e., groups of traits representing function types - diet, temperature preference and morphology) or *a posteriori*, forming trait combinations that maximized or minimized functional dispersion patterns (SES) using our forward selection procedure (see Methods). Figure 2 presents the distribution of SES across all lakes, comparing the two combination methods and considering all traits combined. The forward procedure, which combines traits regardless of their *a priori* functional types, improved the detection of functional dispersion patterns (Figure 2). The combination of all traits resulted in weaker dispersion, confirming that under- or overdispersed trait combinations can lead to seemingly random species assemblages. Strength and directionality (i.e., average SES across lakes) of community trait patterns depended on the species pool (spatial scale) used to estimate SES (Figure 2). While diet-related traits consistently maximized overdispersion across all spatial scales, temperature preference appeared underdispersed at the primary watershed but overdispersed at the secondary watershed level. At the provincial level, the average SES for temperature-related traits was close to zero, demonstrating no major directionality in functional dispersion. This suggests that different selection mechanisms operate at these scales in selecting species based on their diet traits.

**Figure 2.**
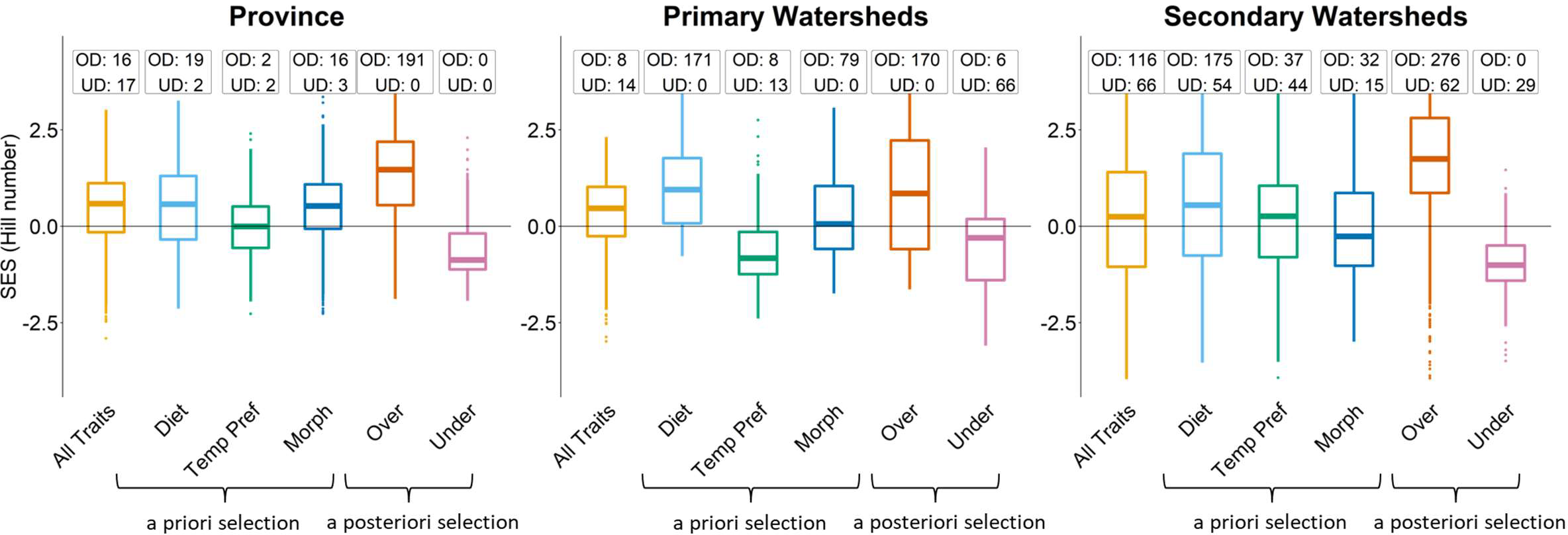
Distributions of SES for different sets of traits: All Traits (all traits pooled together), Morph (Morphology Traits), Temp Pref (Temperature Preference), Under (Traits optimizing under-dispersion), Over (Traits optimizing over-dispersion). The SES are calculated with different species pools defined with geographic coordinates (1 Province pool, 3 Primary Watersheds, 21 Secondary Watersheds). In the text boxes are the number of communities that were significantly overdispersed (OD) and underdispersed (UD). The threshold to determine significance was set at 0.05.

The combinations of traits selected to minimize (underdispersion) or maximize (overdispersion) SES varied across watershed. Specifically, as the spatial scale of species pools decreased (i.e., transitioning from province to primary to secondary watersheds), the combinations of traits associated with the selection process became more diverse: at the secondary watersheds level, the number of unique combination of traits was 14 for underdispersion and 15 for overdispersion (Figure 3). Some traits were found to optimise both patterns, such as Prepectoral Length and Body Depth, while others were only selected to maximise overdispersion, such as Trophic levels, or to minimise underdispersion, like Aspect Ratio.

**Figure 3.**
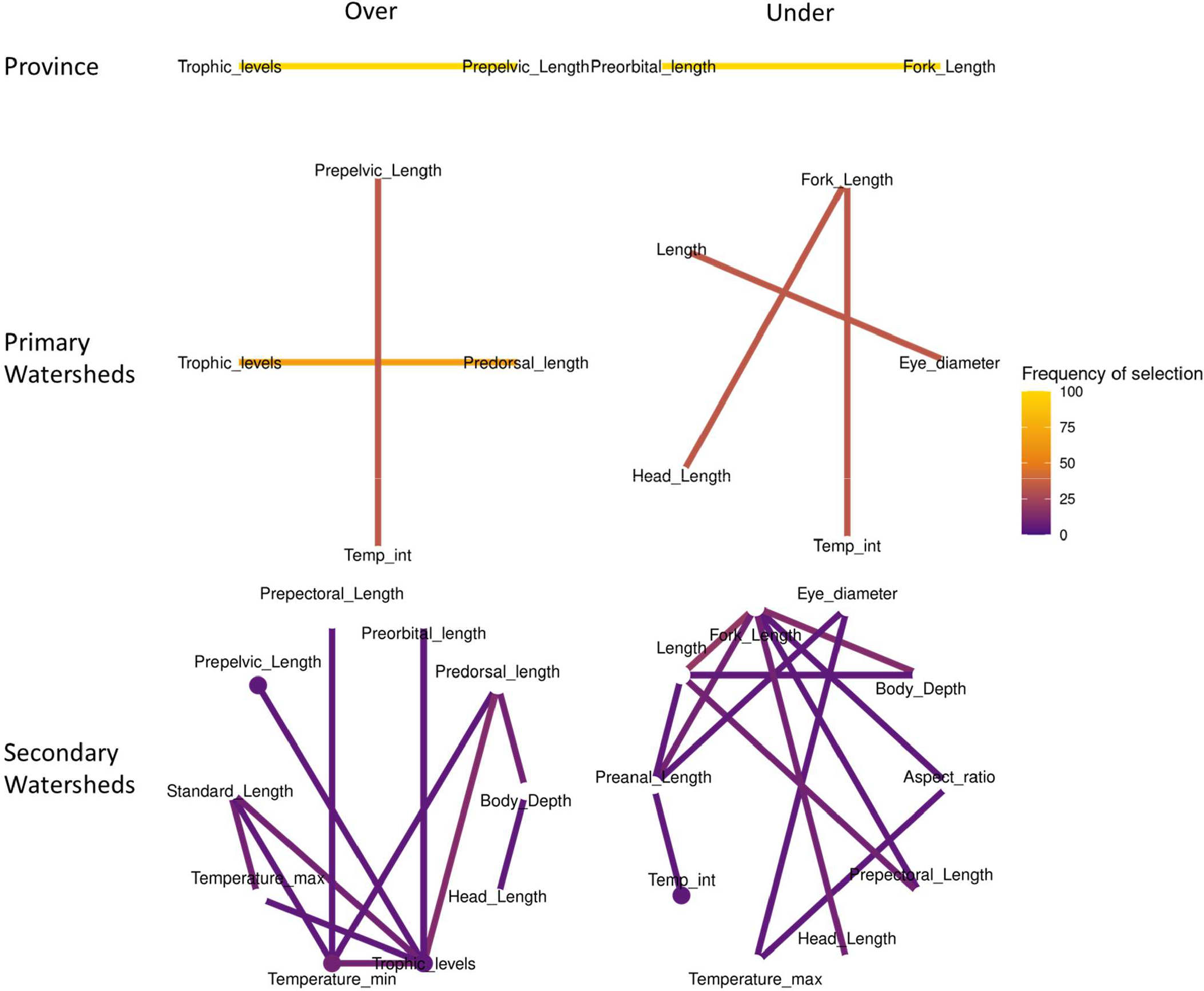
Frequency of the combination of traits selected to optimize either over-dispersion or under-dispersion. We used a stepwise selection method to select traits that would optimize under and over-dispersion for different species pools. The frequency was calculated by counting the number of times a trait was selected and divided by the number of species pools at a given scale (1 Province, 3 Primary Watersheds, 21 Secondary Watersheds). The color of the edge represents the frequency of the selection of the pair: purple means that the pair was not frequently selected together, and yellow means the frequency of selection was high. Some traits were selected on their own, we colored the nodes with the same colour scheme as the edges.

We have also evaluated the variation in selected functional dispersion traits against the patterns of functional dispersion in non-selected traits. Most trait correlations showed weak patterns in community dispersion (Figure 4; see variation in mean absolute correlation across watersheds). However, for some species pools at the secondary watershed scale, certain correlations among traits were relatively high. For those, the network of trait dispersion correlations can potentially identify which traits and modules (i.e., morphology, diet, and temperature) are most correlated and be used to provide further insights into the filtering processes.

**Figure 4.**
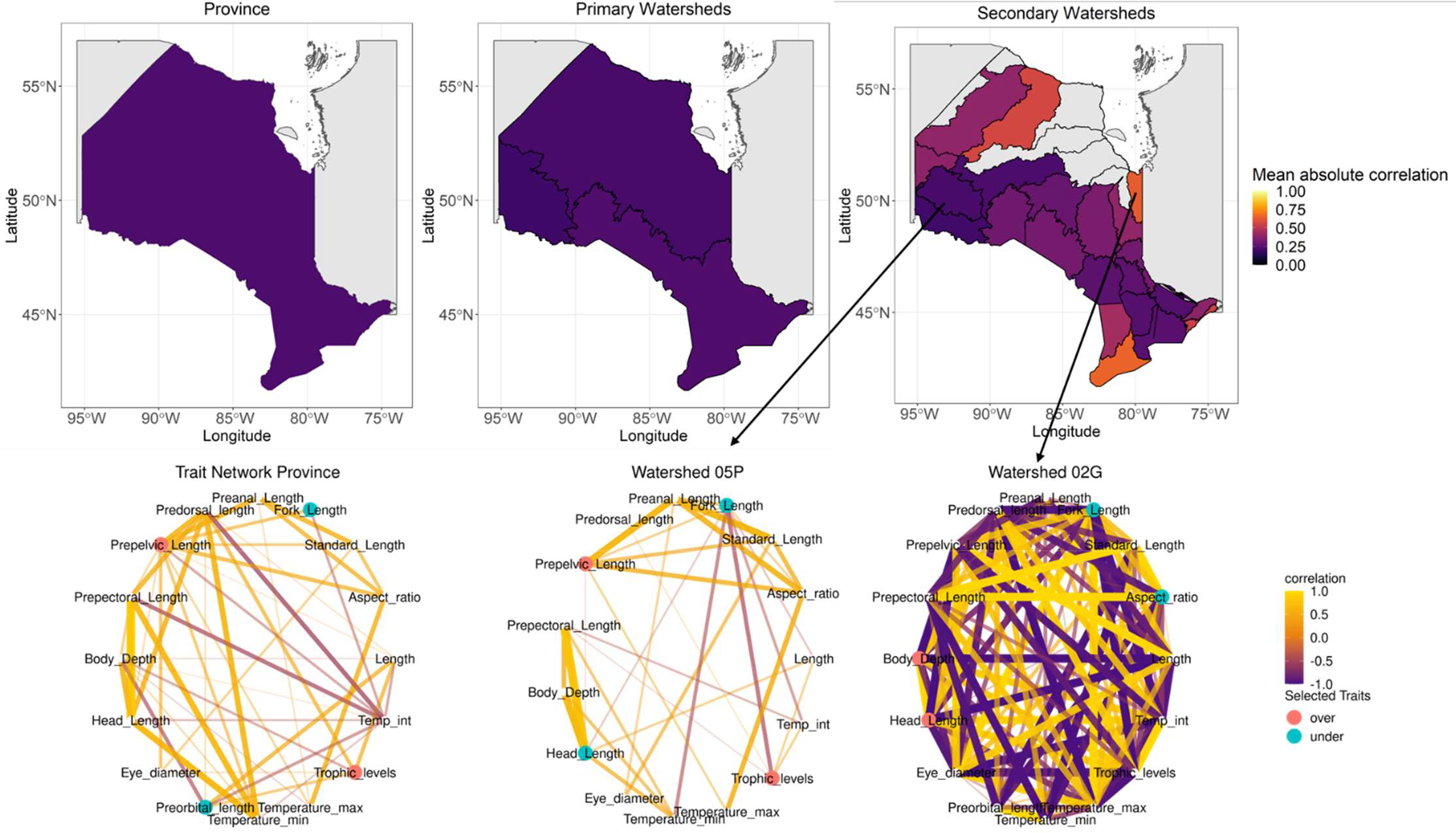
Mean absolute correlation of Standardized Effect Size for single traits calculated for different species pools contained in watersheds. Low mean absolute correlation means that single traits do not have correlated functional dispersion patterns. High mean absolute correlation means that all functional dispersion patterns of single traits are highly correlated. The network represents the Pearson’s correlations among the traits in a specific species pool. Positive correlations (yellow) means that the two traits share similar dispersion patterns: when one is over-dispersed, the other is also over-dispersed. Negative correlations (purple) means that two traits have opposite dispersion patterns: when one is over-dispersed, the other one is under-dispersed. The selected traits are highlighted (in blue, traits selected to optimize under-dispersion. In red, traits selected to optimize over-dispersion). We chose to present the networks for the province and for the most extreme networks possible (Watershed 05P has the weakest correlation network and Watershed 02G the strongest correlation network).

### Functional dispersion patterns: little deviation from null communities

Despite the presence of under and overdispersion patterns in trait combinations (Figure 2), most lake-fish communities were close to what would be expected by random species assemblages from their respective pool (i.e., chance alone). This was particularly true for the *a priori* trait selection (Figure 2). With our null model, we identified statistically significant communities. When all traits were pooled together, the extreme (“significant”) SES values showed equal levels of under and overdispersion (Figure 2). Diet traits were associated with more overdispersed communities, while morphology and temperature preference traits exhibited minor deviations from the null model, with a balance between underdispersion and overdispersion. Our posteriori selection method indicated an overall trend of greater overdispersion in lakes compared to underdispersion, with more extreme SES (significant) values observed in trait combinations maximizing overdispersion (Figure 2).

### Latitudinal and environmental trends in functional dispersion

Although the SES of functional dispersion is often considered to have limited statistical power in detecting community structure, it is commonly observed that its variation among local communities can be strongly linked to environmental variation. In our study, despite the weak signals of community structure based solely on functional dispersion (SES; Figures 2 and 4), we observed pronounced latitudinal (Figure 5) and environmental variation (Figure 6). The latitudinal trends were influenced more by the specific trait combinations used to calculate functional dispersion rather than the species pools in the null communities. Regardless of the species pool employed in the null model, the sign of the trend remained consistent, with a positive trend consistently observed, although the significance of the trend varied.

**Figure 5.**
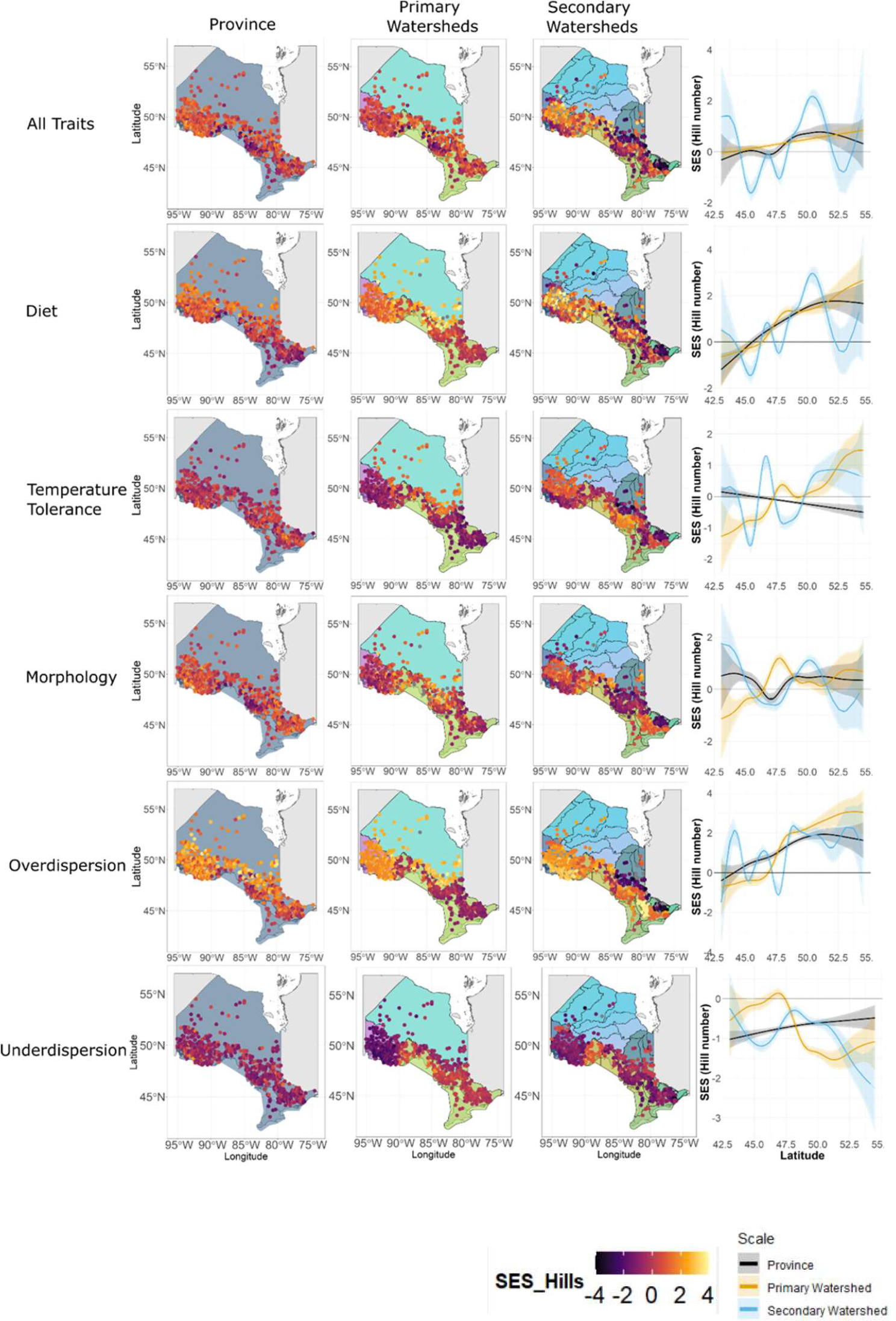
Latitudinal gradients of Standard Effect Size for the different sets of traits and species pools in lake-fish communities of Ontario, Canada. The latitudinal trends were obtained using Generalized Additive Models (GAMs).

**Figure 6.**
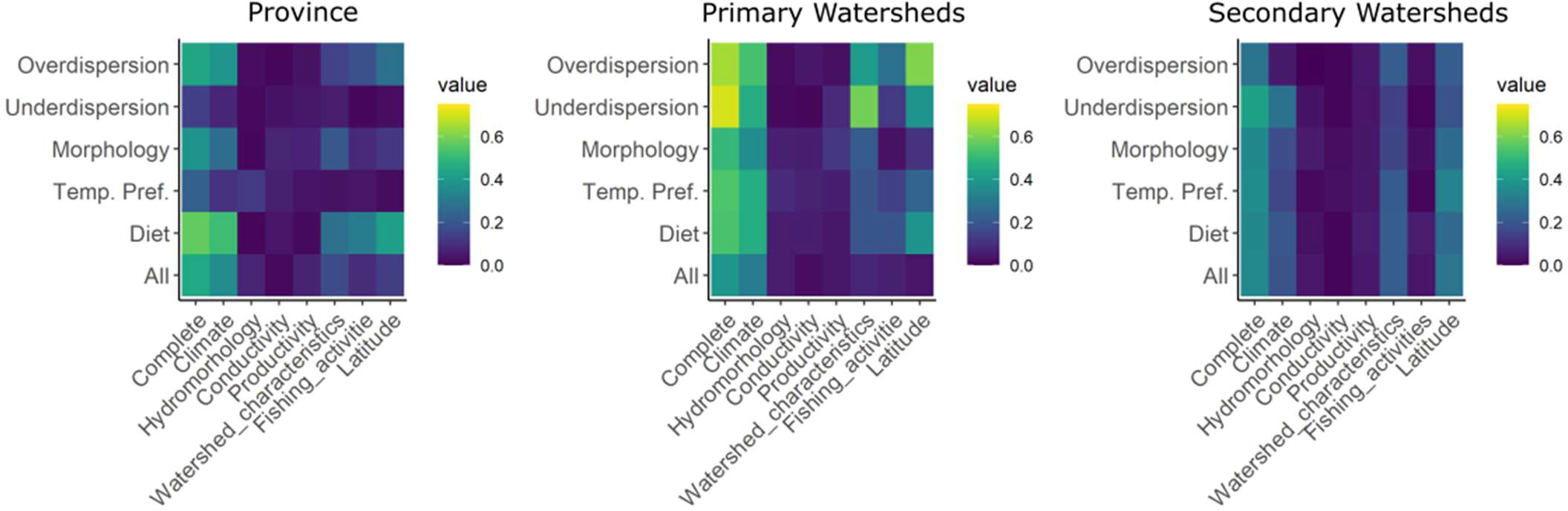
Performances of different environmental models for the SES for all trait sets. We ran a Generalized Additive Models (GAMs) model with all the environmental categories (Complete), for each category of environmental variables, latitude, and for each trait sets. Values represent adjusted GAM R^2^ as an index of model performance (going from 0 to 0.75).

The SES for each set of traits and species pool followed non-linear latitudinal trends (Figure 5). Functional dispersion for temperature preference traits appeared to have a negative latitudinal trend at the Provincial level: southern lakes were more over-dispersed and northern lakes are more underdispersed. However, this trend seemed to reverse when species pools were spatially constrained (i.e., Primary and Secondary Watershed levels). Conversely, the patterns in functional dispersion for diet traits were positive: southern lakes were under-dispersed, whereas northern lakes were over-dispersed. This trend was very strong at the Primary watershed scale. For morphology traits, and when all traits were pooled together, no latitudinal trend was observed: SES oscillated around 0 across the entire latitudinal gradient. For the overdispersion combination of traits, there was a positive latitudinal at all scales, and for the underdispersion set, there was a slightly positive trend at the provincial level but a highly negative latitudinal trend at Primary and Secondary Watershed scales.

The performance of generalized additive models (GAMs) linking trait dispersion patterns (SES) and environmental variation varied between trait combinations and environmental categories (Figure 6). Climate conditions explained the highest proportion of variation in SES, particularly at the Primary Watershed scale. Watershed characteristics accounted for a small amount of the variation in the diet set at the Provincial scale, and a larger amount of variation in overdispersion and underdispersion sets at the Primary Watershed scale. At the Secondary Watershed scale, the performance of the different models was weaker (Figure 6). Importantly, while the predictive capacity (R^2^) changed with spatial scale, the relative performance of the models according to a given environmental category changed little. This suggests that, in our study system, the combination of traits is more important than the spatial scale (species pool).

## DISCUSSION

Using a comprehensive database spanning almost 700 lake-fish communities, we determined that detecting trends in functional dispersion can heavily rely on the combination of traits while estimating trait dispersion patterns. When all traits were pooled together, the geographical and environmental trends in our study system were less apparent, while defining trait sets based on specific combinations revealed clearer geographical trends. These findings are consistent with previous studies (Trisos et al. 2014; Lopez et al. 2016; Côte et al. 2019) indicating that indiscriminate pooling of traits obscures signals related both to under- and overdispersion. Moreover, we demonstrated that combining traits with different functions decreases our understanding of how the environment acts in selecting species to compose local communities as a function of their traits. Underdispersion, typically associated with environmental filtering, was more pronounced when defining species pools at broader spatial scales, while overdispersion, indicating negative species interactions, was predominantly observed when species pools were smaller (Figure 3). These results are consistent with the established notion that environmental filtering (e.g., physiological constraints) plays a greater role at large spatial scales, whereas species interactions become more important at smaller scales (Lessard et al. 2015; Bañares-de-Dios et al. 2020).

### Trait selection: Pooling a priori versus selecting a posteriori

We contrasted two trait selection methods: one based on fish biology knowledge and the other optimizing trait combinations for over- and underdispersion. The first method considers *a priori* hypotheses underlying trait functions but may weaken community trait patterns when pooling non-relevant traits that respond differently across environmental gradients. Conversely, the second method (*a posteriori*) improves predictions of local species composition based on trait combinations but may be more complex to interpret functionally. Initially, using our *a posteriori* trait pooling, functional dispersion of most lake fish communities still did not differ from what was expected by a null model except for Diet sets. This indicates a lack of strong signal of environmental filtering and only weak signals of competition non-relevant traits. These results are consistent with a previous study on Ontario lake-fish communities using datasets from 1980, where 26 pooled traits revealed overdispersion in only 376 of 6977 communities and only 134 were underdispersed (Lamothe et al. 2018). The limited dispersion patterns for most communities could be due to conflicting patterns (over-versus underdispersion) being diluted from the pooling of numerous traits. Alternatively, it is possible that processes other than environmental filtering and competition primarily drive community assembly for Ontario’s freshwater fish communities. Other processes, such as stochastic processes or predation, can significantly structure communities and do not produce specific patterns in functional dispersion (Adler et al. 2013).

For *a priori* sets of traits, we expected Diet to be mainly overdispersed and conversely, Morphology and Temperature Preference to be underdispersed (see Introduction). Only diet clearly followed this expectation. Morphology traits were on average overdispersed at the Province and Primary Watershed scales, and on average close to random at the Secondary watershed scale. Temperature Preference was, on average, close to what was expected by chance alone, with as many overdispersed as underdispersed (lake) communities. This pattern could be explained either by pooling traits with different functional dispersion that cancel each other out, or by an even distribution of thermal environments within lakes that favour a diversity of temperature preferences in the communities.

For *a posteriori* trait combinations, we expected diet traits to be more often selected to maximise overdispersion, whereas traits related to morphology and temperature preference to be selected to maximize underdispersion. This was only partially observed. For instance, trophic levels (which are associated with diet) were selected to maximize overdispersion at the Provincial level and for most Primary and Secondary Watersheds. Traits that would be selected to minimise underdispersion were often from the Morphology dataset (e.g., Body Depth and Aspect Ratio), suggesting that those traits are sensitive to environmental filtering. However, two morphology traits were often selected to maximize overdispersion (Predorsal Length and Prepelvic Length). This demonstrates that when morphology traits are pooled together in one set, they can weaken the ability to detect functional dispersion.

Although we observed strong patterns of overdispersion for the sets designed to maximize overdispersion, we did not observe corresponding strong patterns of underdispersion, even when explicitly maximizing it. Lamothe et al. (2018) made a similar finding in which overdispersion was more often detected than underdispersion. This suggests that, in our study system, competition may be a stronger force in structuring communities than environmental filtering. One possible explanation is that these northern fish communities are well adapted to more extreme environments, lowering the importance of filtering relative to competition. Overdispersion was also easier to detect at smaller scales, aligning with general theory that competitive interactions are more important at finer scales (here spatially constrained species pools) (Lessard et al. 2012; Bañares-de-Dios et al. 2020).

Generally, correlations between individual traits were weakly correlated, suggesting different possibilities regarding their relationships with the assembly process. One possibility is that traits are largely independent, with only a select few playing significant roles in the assembly process. Alternatively, responses of traits to environmental gradients maybe distinct from one another, leading to weak correlations. Finally, the weak correlation in dispersion patterns among traits may be attributed to assembly processes that are more complex than can be unraveled by current metrics of functional dispersion. Our trait selection procedure either maximized overdispersion or minimized underdispersion according to a particular species pool. It is possible, however, that complex patterns leading to weak correlation among dispersion patterns are related to local rather than regional selection of traits. In this case, a procedure that selects traits to increase (or decrease) dispersion for each lake separately could be an avenue to uncover the more complex ways in which traits are involved in community assembly.

### Geographical dispersion patterns: complex interactions between traits and abiotic conditions

We anticipated that temperature preference and Morphology traits would be underdispersed (negative SES) due to their sensitivity to environmental filtering, while Diet traits would be overdispersed (positive SES) due to competitive interactions. We also hypothesized stronger environmental filtering for Temperature preference traits and increased competition related to Diet traits in northern lakes. However, the results did not align with these expectations, especially for simple variation in SES values (i.e., independent of latitudinal and environmental predictors). Across all trait sets, the medians SES values were close to zero, indicating equal occurrences of over- and underdispersion. Even traits expected to be sensitive to environmental filtering displayed overdispersion, while traits supposed to be linked to competition showed underdispersion. Either Ontario freshwater fish communities are mostly randomly assembled communities, resulting in weak SES values or traits may be sensitive to multiple filters simultaneously. For example, Côte et al. (2019) discussed how traits like fish size can be linked to both environmental requirements and competitive ability. The fact that our environmental models of SES resulted in strong predictive ability, indicates that trait-related assembly processes in these lakes are indeed complex and not random.

For Temperature preference traits, communities were strongly overdispersed in southern lakes and underdispersed in northern lakes. The overdispersion of Temperature preference traits in southern lakes could be potentially explained by environmental heterogeneity. Local heterogeneity may lead to overdispersion when underdispersion is expected, as environmental filters may apply at a finer scale than how communities are defined (Adler et al. 2013; Lopez et al. 2016; e.g., microgeographic variation within lakes).

The models that related climate variables to functional patterns performed rather well, particularly for Diet traits. For Ontario fishes, climate is one of the strongest constraints for successfully colonizing lakes (Crossman & Mandrak 1992; Jackson et al. 2001; Alofs & Jackson 2015). The strong latitudinal trends observed are highly influenced by the significant climate gradient present in Ontario. Fishing activities partially explained the variation in dispersion patterns across communities. Fishing activities are known to meaningfully impact functional diversity by, for instance, targeting species with similar functional traits (Olden et al. 2007). The Watershed characteristics explained a part of the variance in the functional dispersion patterns of lake-fish communities in Ontario: historical context also impacts community assembly by reflecting the post-glaciation colonization as well as contemporary and historical connectivity (Layeghifard et al. 2015; Loewen et al. 2022).

### The importance of the scale in detecting functional patterns

The influence of spatial scales was evident in determining the optimal combination of traits leading to under- or overdispersion. Certain traits, such as trophic level, were specifically selected to minimize underdispersion at larger scales while maximizing overdispersion at smaller scales. Trophic level is directly linked to resource exploitation in fish and is expected to be responsive to competition. Its selection to optimize both patterns at different scales highlights the significant impact scales can have on our ability to detect underlying mechanisms. As discussed, competition is generally easier to detect at more local scales (Götzenberger et al. 2012; De Bello et al. 2013). For example, Bañares-de-Dios et al. (2020) demonstrated that defining null communities using larger spatial scales maximizes trait diversity in species pools. Consequently, environmental filtering becomes the dominant mechanism explaining trait variation among communities. In contrast, at smaller spatial scales, the species pools exhibit a reduced diversity of traits, making competitive exclusion more readily apparent.

The *a posteriori* method also highlighted the importance of local context in determining how different processes could shape functional diversity in Ontario’s lake-fish communities. At more local scales, the diversity of possible combinations that would maximise or minimise functional dispersion was greater than at broader scales. In addition, some traits could be selected to maximise one pattern in one secondary watershed but could be selected to optimise the other pattern in another watershed. These findings indicate that as the spatial scale decreases, the complexity of local processes influencing the selection of species in lake communities increases.

## CONCLUSION

Our study demonstrates the importance of trait selections for detecting and interpreting patterns of functional dispersion in ecological communities. Pooling traits together obscures the unique responses of individual traits to environmental gradients and hampers our ability to understand how communities are assembled. By separating traits, we gained a more refined understanding of the complex processes underlying Ontario lake-fish communities. Our key findings revealed that traits with distinct *a priori* functional expectations exhibit complex relationships that influence the assembly process. Finally, we have outlined future selection procedures that can improve further our ability to undercover even more complex patterns of species associations based on traits.

## Acknowledgements

The data presented in this article were sampled and curated by the Ontario Ministry of Natural Resources and Forestry, we thank Dr. Cindy Chu for graciously giving us access to the Ontario Broadscale Monitoring Program data. This work was funded by FONCER-BIOS2 grant. We thank the members of the Quantitative and Community Ecology Laboratory and Fraser Laboratory for their feedbacks during the development of this project and their friendly reviews of the different drafts of this manuscript.

## Conflict of Interests

The authors do not have any conflict of interest.

## Authors contribution

The study was designed by Alexandra Engler, Dylan Fraser, and Pedro Peres-Neto. The analysis and interpretation of the results were mainly conducted by Alexandra Engler with critical inputs from Dylan Fraser and Pedro Peres-Neto. Alexandra Engler led the writing of the drafts and all the authors have edited and approved the submitted draft.

## Notes

https://doi.org/10.5281/zenodo.8038809

### Competing Interest Statement

The authors have declared no competing interest.

## REFERENCES

Adler PB, Fajardo A, Kleinhesselink AR, Kraft NJB. 2013. Trait-based tests of coexistence mechanisms. Ecol Lett. 16(10):1294–1306.

Alofs KM, Jackson DA. 2015. The abiotic and biotic factors limiting establishment of predatory fishes at their expanding northern range boundaries in Ontario, Canada. Glob Chang Biol. 21(6):2227–2237.

Appleberg M. 2000. Swedish standard methods for sampling freshwater fish with multi-mesh gillnets : stratified random sampling with Nordic multi-mesh gillnets provide reliable whole-lake estimates of the relative abundance and biomass of freshwater temperate lakes. Fisk Inf (Sweden).(2000:1):31.

Bañares-de-Dios G, Macía MJ, Granzow-de la Cerda í, Arnelas I, Martins de Carvalho G, Espinosa CI, Salinas N, Swenson NG, Cayuela L. 2020. Linking patterns and processes of tree community assembly across spatial scales in tropical montane forests. Ecology. 101(7):1–13.

De Bello F, Vandewalle M, Reitalu T, Lepš J, Prentice HC, Lavorel S, Sykes MT. 2013. Evidence for scale- and disturbance-dependent trait assembly patterns in dry semi-natural grasslands. J Ecol. 101(5):1237–1244.

Cadotte MW, Tucker CM. 2017. Should Environmental Filtering be Abandoned? Trends Ecol Evol. 32(6):429–437.

Chiu CH, Chao A. 2014. Distance-based functional diversity measures and their decomposition: A framework based on hill numbers. PLoS One. 9(7).

Cornwell WK, Ackerly DD. 2009. Community assembly and shifts in plant trait distributions across an environmental gradient in coastal California. Ecol Monogr. 79(1):109–126.

Côte J, Kuczynski L, Grenouillet G. 2019. Spatial patterns and determinants of trait dispersion in freshwater fish assemblages across Europe. Glob Ecol Biogeogr. 28(6):826–838.

Crossman E., Mandrak NE. 1992. Postglacial dispersal of freshwater fishes into Ontario. Can J Zool. 70(11):2247–2259.

Froese R, Pauly D. 2019. FishBase,version (12/2019) . http://www.fishbase.org

Götzenberger L, de Bello F, Bråthen KA, Davison J, Dubuis A, Guisan A, Lepš J, Lindborg R, Moora M, Pärtel M, et al. 2012. Ecological assembly rules in plant communities-approaches, patterns and prospects. Biol Rev. 87(1):111–127.

Gower JC. 1971. A General Coefficient of Similarity and Some of Its Properties. Int Biometric Soc. 27(4):857–871.

Henriques-Silva R, Kubisch A, Peres-Neto PR. 2019. Latitudinal-diversity gradients can be shaped by biotic processes: new insights from an eco-evolutionary model. Ecography (Cop). 42(2):259–271.

Jackson DA, Peres-Neto PR, Olden JD. 2001. What controls who is where in freshwater fish communities – the roles of biotic, abiotic, and spatial factors. Can J Fish Aquat Sci. 58(1):157–170.

Kneitel JM, Chase JM. 2004. Trade-offs in community ecology: Linking spatial scales and species coexistence. Ecol Lett. 7(1):69–80.

Laliberté E, Legendre P, Shipley B. 2014. FD: measuring functional diversity from multiple traits, and other tools for functional ecology. R package version 1.0-12.

Lamothe KA, Alofs KM, Jackson DA, Somers KM. 2018. Functional diversity and redundancy of freshwater fish communities across biogeographic and environmental gradients. Divers Distrib. 24(11):1612–1626.

Layeghifard M, Makarenkov V, Peres-Neto PR. 2015. Spatial and species compositional networks for inferring connectivity patterns in ecological communities. Glob Ecol Biogeogr. 24(6):718–727.

Lessard J-P, Belmaker J, Myers JA, Chase JM, Rahbek C. 2012. Inferring local ecological processes amid species pool influences. Trends Ecol Evol. 27(11):600–607.

Lessard JP, Weinstein BG, Borregaard MK, Marske KA, Martin DR, McGuire JA, Parra JL, Rahbek C, Graham CH. 2015. Process-based species pools reveal the hidden signature of biotic interactions amid the influence of temperature filtering. Am Nat. 187(1):75–88.

Lester NP, Sandstrom S, de Kerchhove DT, Armstrong K, Ball H, Amos J, Dunkley T, Rawson M, Addison P, Dextrase A, et al. 2020. Standardized Broad - Scale Management and Monitoring of Inland Lake Recreational Fisheries : An Overview of the Ontario Experience. Fisheries. 46(3):107–118.

Loewen CJG, Jackson DA, Chu C, Alofs KM, Hansen GJA, Honsey AE, Minns CK, Wehrly KE. 2022. Bioregions are predominantly climatic for fishes of northern lakes. Glob Ecol Biogeogr. 31(2):233–246.

Lopez BE, Burgio KR, Carlucci MB, Palmquist KA, Parada A, Weinberger VP, Hurlbert AH. 2016. A new framework for inferring community assembly processes using phylogenetic information, relevant traits and environmental gradients. One Ecosyst. 1:1–24.

Magneville C, Loiseau N, Albouy C, Casajus N, Claverie T, Escalas A, Leprieur F, Maire E, Mouillot D, Villéger S. 2022. mFD: an R package to compute and illustrate the multiple facets of functional diversity. Ecography (Cop). 2022(1):1–15.

Magnuson JJ, Tonn WM, Banerjee A, Toivonen J, Sanchez O, Rask M. 1998. Isolation vs. extinction in the assembly of fishes in small northern lakes. Ecology. 79(8):2941–2956.

Mammola S, Carmona CP, Guillerme T, Cardoso P. 2021. Concepts and applications in functional diversity. Funct Ecol. 35(9):1869–1885.

Mason NWH, De Bello F, Mouillot D, Pavoine S, Dray S. 2013. A guide for using functional diversity indices to reveal changes in assembly processes along ecological gradients. J Veg Sci. 24(5):794–806.

Mayfield MM, Levine JM. 2010. Opposing effects of competitive exclusion on the phylogenetic structure of communities. Ecol Lett. 13(9):1085–1093.

McGill BJ, Enquist BJ, Weiher E, Westoby M. 2006. Rebuilding community ecology from functional traits. Trends Ecol Evol. 21(4):178–185.

Mouillot D, Dumay O, Tomasini JA. 2007. Limiting similarity, niche filtering and functional diversity in coastal lagoon fish communities. Estuar Coast Shelf Sci. 71(3–4):443–456.

Mouillot D, Loiseau N, Grenié M, Algar AC, Allegra M, Cadotte MW, Casajus N, Denelle P, Guéguen M, Maire A, et al. 2021. The dimensionality and structure of species trait spaces. Ecol Lett. 24(9):1988–2009.

Münkemüller T, de Bello F, Meynard CN, Gravel D, Lavergne S, Mouillot D, Mouquet N, Thuiller W. 2012. From diversity indices to community assembly processes: A test with simulated data. Ecography (Cop). 35(5):468–480.

Munoz F, Klausmeier CA, Gaüzère P, Kandlikar G, Litchman E, Mouquet N, Ostling A, Thuiller W, Algar AC, Auber A, et al. 2023. The ecological causes of functional distinctiveness in communities. Ecol Lett.(October):1–2.

Olden JD, Hogan ZS, Zanden MJ Vander. 2007. Small fish, big fish, red fish, blue fish: Size-biased extinction risk of the world’s freshwater and marine fishes. Glob Ecol Biogeogr. 16(6):694–701.

Olden JD, Jackson DA, Peres-Neto PR. 2001. Spatial isolation and fish communities in drainage lakes. Oecologia. 127(4):572–585.

Peterson, Ryan A. 2021. Finding Optimal Normalizing Transformations via bestNormalize. R J . 13(1):310. https://petersonr.github.io/bestNormalize/

R Core Team. 2020. R: A language and environment for statistical computing. https://www.r-project.org/

Schleuter D, Daufresne M, Veslot J, Mason NWH, Lanoiselée C, Brosse S, Beauchard O, Argillier C. 2012. Geographic isolation and climate govern the functional diversity of native fish communities in European drainage basins. Glob Ecol Biogeogr. 21(11):1083–1095.

Trisos CH, Petchey OL, Tobias JA. 2014. Unraveling the interplay of community assembly processes acting on multiple niche axes across spatial scales. Am Nat. 184(5):593–608.

Villéger S, Brosse S, Mouchet M, Mouillot D, Vanni MJ. 2017. Functional ecology of fish: current approaches and future challenges. Aquat Sci. 79(4):783–801.

Wood SN, Pya N, Säfken B. 2016. Smoothing Parameter and Model Selection for General Smooth Models. J Am Stat Assoc. 111(516):1548–1563.

Zhu L, Fu B, Zhu H, Wang C, Jiao L, Zhou J. 2017. Trait choice profoundly affected the ecological conclusions drawn from functional diversity measures. Sci Rep. 7(1):1–13.

