## Supplemental Materials for "Vive la difference: why, how, and what trait combinations improve functional community ecology"

**APPENDIX 1: Supplementary tables and figures**

**Talbe S1:** Detailed descriptions of the traits and their units (extracted from FishBase (Froese & Pauly 2019))

| Trait | Unit | Meaning |
| --- | --- | --- |
| Aspect Ratio | NA | Caudal height/Caudal fin area |
| Body Depth | Percentage of Total Length | Height of the body/Length |
| Eye Diameter | Percentage of Head Length | Eye diameter/Length of the head |
| Fork Length | Percentage of Total Length | Proportion of length to the fork over total length |
| Head Length | Percentage of Total Length |  |
| Length | Cm | Common length |
| Maximum Temperature | Celsius | Maximum Temperature in the species range |
| Minimum Temperature | Celsius | Minimum Temperature in the species range |
| Preanal Length | Percentage of Total Length | Length between the mouth to pre-anal fin/Length |
| Predorsal Length | Percentage of Total Length | Length between the mouth to the pre-dorsal fin /Length |
| Preorbital Length | Percentage of Head Length | Length between the mouth to eye /Length |
| Prepectoral Length | Percentage of Total Length | Length between the mouth to pre-pectoral fin/Length |
| Prepelvic Length | Percentage of Total Length | Length between the mouth to pre-pelvic fin/Length |
| Standard Length | Percentage of Total Length | Length to the tail |
| Temperature Interval | Celsius | Minimum Temperature – Maximum Temperature |

**Table S2:** Description of environmental variables, their units and the transformation method used to normalise the data.

| Name | Description | Category | | Unit | Transformation |
| --- | --- | --- | --- | --- | --- |
| AirTemp_8110 | Mean Annual Air Temperature for 1981-2010 | Climate | | Celsius | Ordered quantiles normalization (orderNorm) |
| Alkalinity.mg.L.CaCO3 | concentration of CaCO3 in the lake | Conductivity | | Mg/L | Log10(x) |
| Alkalinity.mg.L.CaCO3_pctl | Percentile of concentration of CaCO3 in the lake class (Lake size) | Conductivity | |  | Ordered quantiles normalization (orderNorm) |
| Altitude_m | Altitude above the sea level of the lake | Watershed characteristic | | m | Ordered quantiles normalization (orderNorm) |
| Amonia_Amonium.mg.L | Concentration of Amonia and Amonium | Productivity | | Mg/L | Ordered quantiles normalization (orderNorm) |
| Amonia_Amonium.mg.L._pctl | Percentiles of the concentration of Amonia and amonium within lake class | Productivity | |  | Ordered quantiles normalization (orderNorm) |
| Angling Pressure | Annual angling pressure (angler-hours/ha-year) - based on aerial survey counts | Fishing activities | angler-hours/ha-year | | Ordered quantiles normalization (orderNorm) |
| Area_km2 | Surface Area of the lake | Hydromorphology | | Km2 | Ordered quantiles normalization (orderNorm) |
| Aut0 | Average date of the last day above 0 from 1981-2010 | Climate | | Days | Ordered quantiles normalization (orderNorm) |
| Calcium.mg.L | Concentration of Calcium | Conductivity | | Mg/L | Ordered quantiles normalization (orderNorm) |
| Calcium.mg.L_pctl | Percentile of Concentration of Calcium | Conductivity | |  | Ordered quantiles normalization (orderNorm) |
| Chloride.mg.L. | Concentration of Chloride | Conductivity | | Mg/L | Ordered quantiles normalization (orderNorm) |
| Chloride.mg.L._pctl | Percentile of the concentration of Chloride within lake size class | Conductivity | |  | Ordered quantiles normalization (orderNorm) |
| Conductivity.uS.cm.s. | Conductivity | Conductivity | | uS/cm/s | Ordered quantiles normalization (orderNorm) |
| Conductivity.uS.cm.s._pctl | Percentiles of the conductivity within lake size class | Conductivity | |  | Box Cox Transformation |
| Conservation_Land | Conservation status (1 implies some form of conservation status) | Fishing activities |  | | Arcsin transformation |
| DD5_8110 | Degree Days above 5C for 1981-2010 | Climate | |  | Ordered quantiles normalization (orderNorm) |
| Depth_Max | Maximum Depth | Hydromorphology | | m | Box Cox Transformation |
| Depth_Mn | Mean Depth | Hydromorpholgy | | m | Log10(x) |
| DIC..mg.L. | Dissolved Inorganic Carbon | Productivity | | Mg/L | Box Cox Transformation |
| DIC..mg.L._pctl | Percentiles of the concentration of DIC within lake class | Productivity | |  | Ordered quantiles normalization (orderNorm) |
| DOC..mg.L_pctl | Percentiles of the concentration of DOC within lake class | Productivity | |  | Ordered quantiles normalization (orderNorm) |
| DOC..mg.L | Dissolved Organic Carbon | Productivity | | Mg/L | Box Cox Transformation |
| FreezDD | Cumulative Degree Days with a temperature <0C | Climate | |  | Ordered quantiles normalization (orderNorm) |
| Hypo.Space.Area.obs | Observed Hypolimnetic Area: Area of the layer of water below the thermocline | Hydromorphology | | Km2 | Ordered quantiles normalization (orderNorm) |
| Hypo.Space.Area.pred | Predicted Hypolimnetic Area: Area of the layer of water below the thermocline | Hydromorphology | | Km2 | Ordered quantiles normalization (orderNorm) |
| Hypo.Space.Vol.pred | Predicted hypolimnetic Volume | Hydromorphology | | M3 | Ordered quantiles normalization (orderNorm) |
| Hypo.Space.Vol.obs | Observed Hypolimnetic volume | Hydromorphology | | M3 | Ordered quantiles normalization (orderNorm) |
| Iron | Concentration of Iron | Conductivity | | Mg/L | Yeo-Johnson Transformation |
| Iron_pctl | Percentiles of the concentration of iron within lake class | Conductivity | |  | Ordered quantiles normalization (orderNorm) |
| Lak_age | Estimated age of the lake | Spatial | | Kyr | Box Cox Transformation |
| Magnesium.mg.L. | Concentration of Magnesium | Conductivity | | mg/L | Ordered quantiles normalization (orderNorm) |
| Magnesium.mg.L._pctl | Percentiles of the concentration of Magnesium within lake class | Conductivity | |  | Ordered quantiles normalization (orderNorm) |
| MAT_8110 | Mean Annual Air Temperature for 1981-2010 | Climate | | Celsius | Ordered quantiles normalization (orderNorm) |
| Max.surface.T | Maximum Surface Temperature | Climate | | Celsius | Arcsin transformation |
| MxMonTP | Maximum Monthly surface Temperature | Climate | | Celsius | Ordered quantiles normalization (orderNorm) |
| MxWatTP | Boosted estimation of the maximum summer water temperature | Climate | | Celsius | Ordered quantiles normalization (orderNorm) |
| Nitrate.Nitrite.ug.L | Concentration of Nitrate and Nitrite | Productivity | | µg/L | Ordered quantiles normalization (orderNorm) |
| Nitrate.Nitrite.ug.L._pctl | Percentiles of the concentration of Nitrate and Nitrite within lake class | Productivity | |  | Ordered quantiles normalization (orderNorm) |
| No..ice.free.days | Estimated umber of icefree days | Climate | | Days | Ordered quantiles normalization (orderNorm) |
| pArea_LE20 | Proportion of Lake area < 20 m in depth | Hydromorphology |  | | Ordered quantiles normalization (orderNorm) |
| pDays.Cold | Proportion of days Cold (between 8-12C) during icefree period | Climate | |  | Ordered quantiles normalization (orderNorm) |
| pDays.Cool | Proportion of days Cool (12 and 22 C) during icefree period | Climate | |  | Ordered quantiles normalization (orderNorm) |
| pDays.Warm | Proportion of warm days (22-26C) during icefree period | Climate | |  | Ordered quantiles normalization (orderNorm) |
| pH | pH | Conductivity | |  | Arcsin transformation |
| pH_pctl | Percentiles of pH within lake class | Conductivity | |  | Ordered quantiles normalization (orderNorm) |
| Perim_km | Perimeter of the lake | Hydromorphology | | km2 | Ordered quantiles normalization (orderNorm) |
| pLittoral | Proportion of littoral | Hydromorphology | |  | Box Cox Transformation |
| PodDD | Cumulative Degree Days with a temperature >0C | Climate | |  | Yeo-Johnson Transformation |
| PosDays | Number of day > 0C | Climate | |  | Ordered quantiles normalization (orderNorm) |
| PosPrecip | Average rainfall from 1981-2010 | Climate | |  | Ordered quantiles normalization (orderNorm) |
| Potassium.mg.L. | Concentration of Potassium | Conductivity | | Mg/L | Ordered quantiles normalization (orderNorm) |
| Potassium.mg.L._pctl | Percentiles of the concentration of Potatium within lake class | Conductivity | |  | Ordered quantiles normalization (orderNorm) |
| Secchi_Su | Secchi depth of the lake in summer | Productivity | | m | Ordered quantiles normalization (orderNorm) |
| Secchi_Sp | Secchi depth of the lake in spring | Productivity | | m | Box Cox Transformation |
| Shoreline_Development_Factor | Shoreline Development Factor | Hydromorphology | |  | Box Cox Transformation |
| Shoreline_km | Total shoreline of lake (include islands and perimter) | Hydromorphology | | Km | Box Cox transformation |
| Silicate..mg.L. | Concentration of silicate | Productivity | | Mg/L | Box Cox transformation |
| Silicate..mg.L._pctl | Percentiles of the concentration of silicate within lake class | Productivity | |  | Ordered quantiles normalization (orderNorm) |
| Sodium.mg.L | Concentration of Sodium | Conductivity | | Mg/L | Ordered quantiles normalization (orderNorm) |
| Sodium.mg.L._pctl | Percentiles of the concentration of sodium within lake class | Conductivity | |  | Ordered quantiles normalization (orderNorm) |
| Spr0 | Average date of the first day above 0 from 1981 - 2010 | Climate | |  | Center + Scale |
| Sulphate.mg.L. | Concentration of sulphate | Conductivity | | Mg/L | Standardised arcsin |
| Sulphate.mg.L._pctl | Percentiles of the concentration of sulphate within lake class | Conductivity | |  | Ordered quantiles normalization (orderNorm) |
| Summer.Shore.Count | Mean Count of shore fishers in summer | Fishing activities | |  | Square-root Transformation |
| Summer.Vessel.Count | Mean count of fishing boats in summer | Fishing activities | |  | Ordered quantiles normalization (orderNorm) |
| TDS | Total Dissolved Solids | Productivity | | Mg/L | Yeo-Johnson Transformation |
| Thermo.Obs | Observed Thermocline Depth | Climate | | m | Square-root transformation |
| Thermo.Pred | Predicted Thermocline Depth | Climate | | m | Ordered quantiles normalization (orderNorm) |
| TKN.ug.L | Total Kjeldahl Nitrogen | Productivity | | µg/L | Square-root transformation |
| TKN.ug.L_pctl | Percentile of Total Kjeldahl Nitrogen | Productivity | |  | Ordered quantiles normalization (orderNorm) |
| Total.Phosphorus.ug.L | Concentration of Phosphorus+ | Productivity | | µg/L | Box Cox Transformation |
| Total.Phosphorus.ug.L_pctl | Percentile of the concentration of Phosphorus within lake size class | Productivity | |  | Ordered quantiles normalization (orderNorm) |
| TP | Total Phosphorus | Productivity | |  | Standardised arcsin |
| True.Colour..TCU. | True Color | Productivity | | TCU | Box Cox Transformation |
| True.Colour..TCU._pctl | Percentile of True Colour within lake class | Productivity | |  | Ordered quantiles normalization (orderNorm) |
| TSI…Avg | Mean of Trophic Status Index (from Phosphorus and Secchi) | Productivity | |  | Arcsin transformation |
| TSI…Phosphorus | Trophic Status Index based on Phosphorus | Productivity | |  | Yeo-Johnson Transformation |
| TSI…Secchi | Trophic Status Index based on Secchi | Productivity | |  | Square-root transformation |
| TWS_age | Age of the Tertiary Watershed | Watershed Characteristics | |  | Ordered quantiles normalization (orderNorm) |
| TWS_area | Tertiary watershed area | Watershed Characteristics | | Km2 | Ordered quantiles normalization (orderNorm) |
| TWS_eleva | Tertiary watershed elevation (m a.s.l.) | Watershed Characteristics | | m | Ordered quantiles normalization (orderNorm) |
| TWS_elevd | Difference between the lowest and highest point in the tertiary watershed | Watershed Characteristics | | m | Ordered quantiles normalization (orderNorm) |
| Volume | area*max depth | Hydromorphology | | M3 | Ordered quantiles normalization (orderNorm) |
| Waterbody_LID | Lake Identifier |  | |  | NA |
| Winter.huts.Counts | Mean count of ice huts in winter | Fishing activities |  | | Square-root transformation |
| Winter.Open.Ice.Counts | Mean Count of open-ice fishers in winter | Fishing activities |  | | Square-root transformation |

**Figure S1:** NMDS for the different sets of traits and different species pools. Each point represents a species that is present in a species pool.

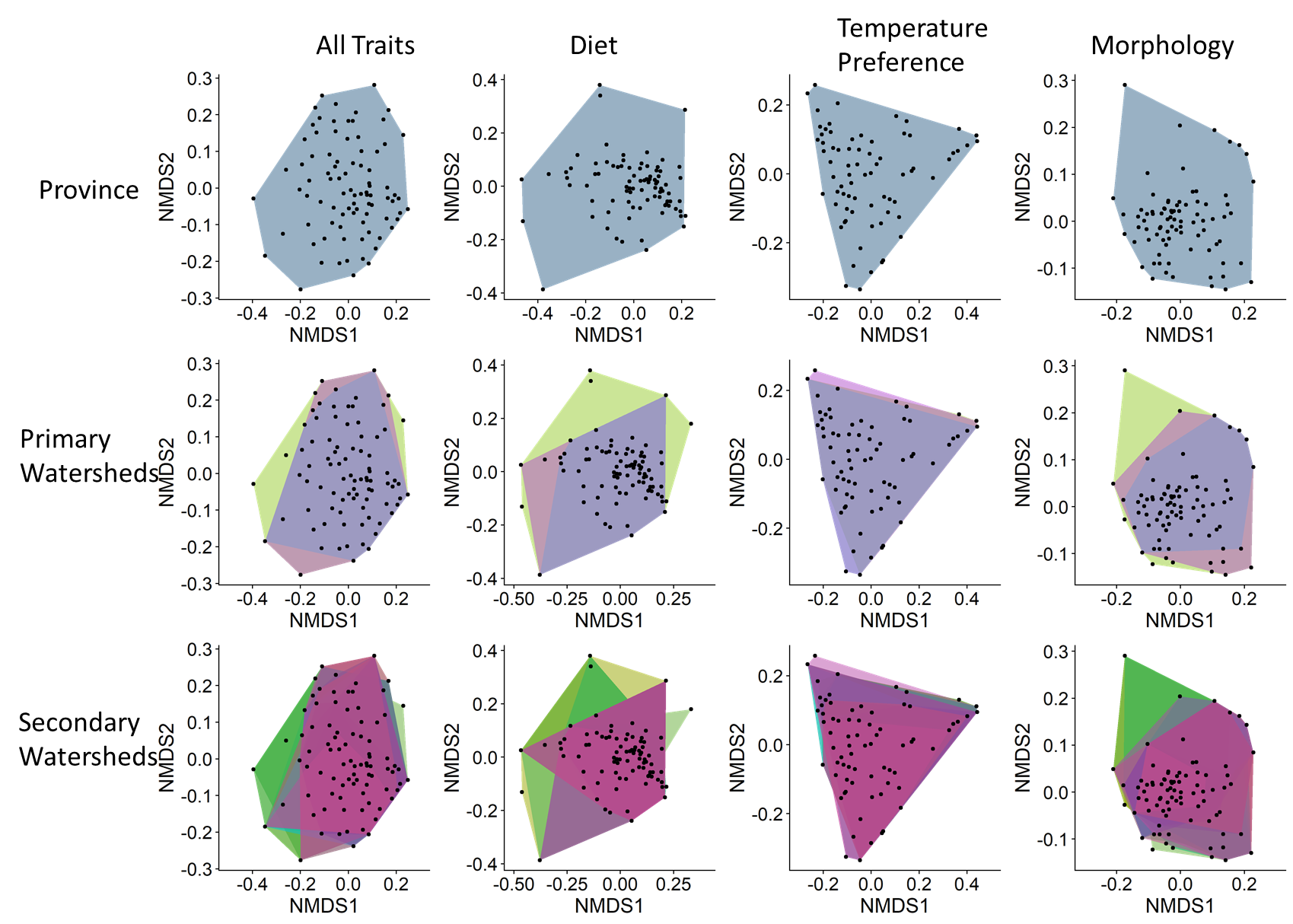

**Figure S2:** Example of networks based on Pearson correlations among the Standard Effect Size calculated for single traits. Positive correlations (yellow) means that the two traits share similar dispersion patterns: when one is over-dispersed, the other is also over-dispersed. Negative correlations (purple) means that two traits have opposite dispersion patterns: when one is over-dispersed, the other one is under-dispersed. The selected traits are highlighted (in blue, traits selected to optimize underdispersion. In red, traits selected to optimize over-dispersion).

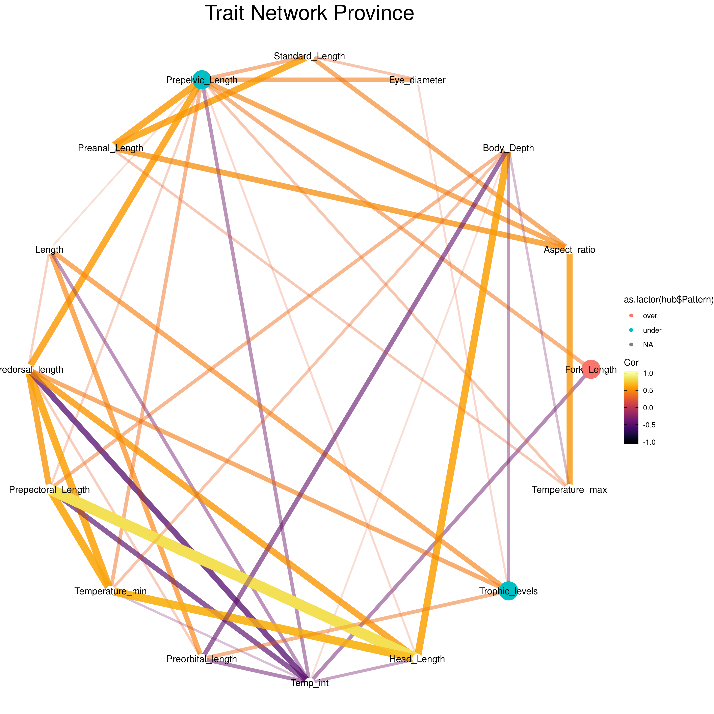

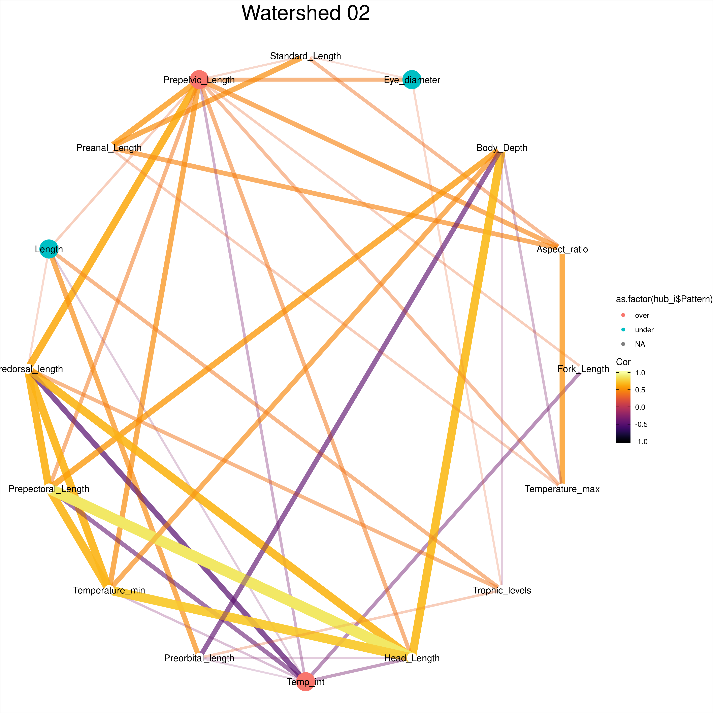

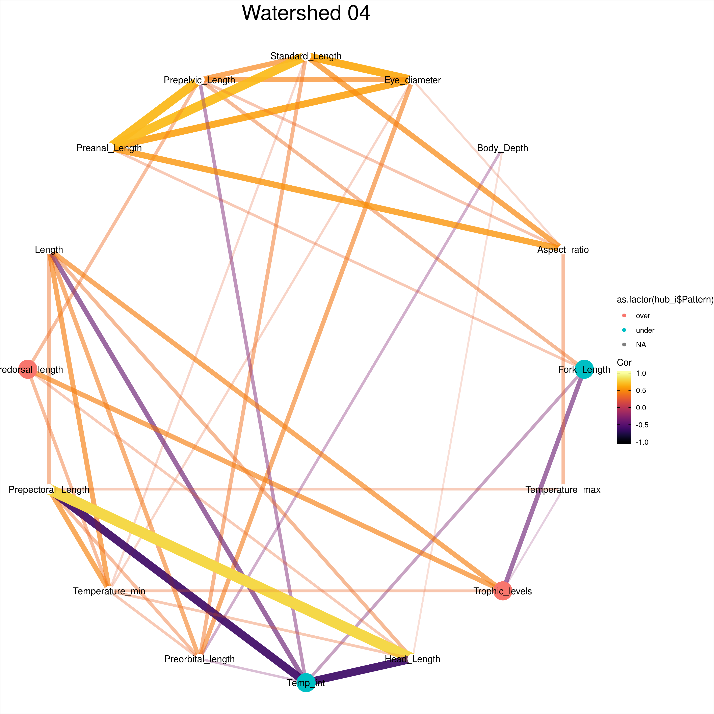

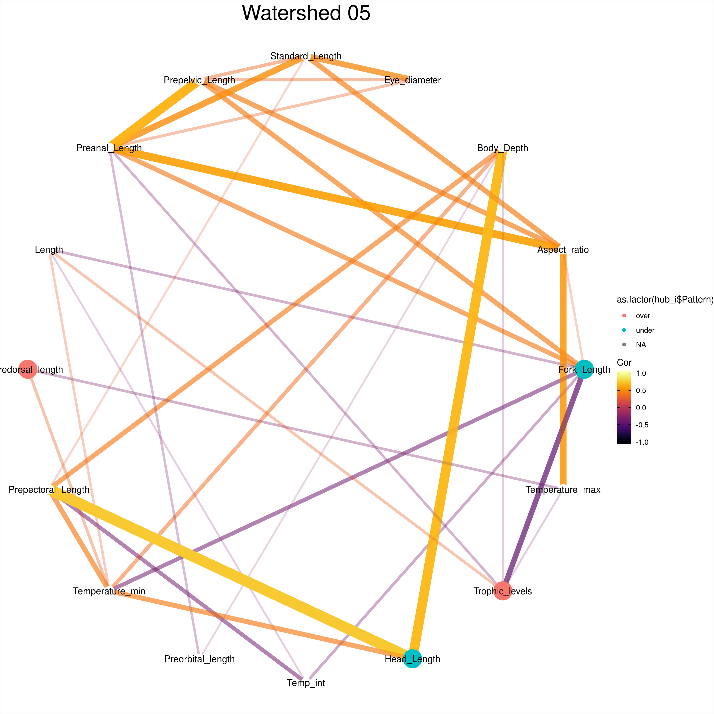

**Figure S3:** Dendrogram of similarities between correlation networks for each species pool. It shows similarities between correlations among the SES of single traits.
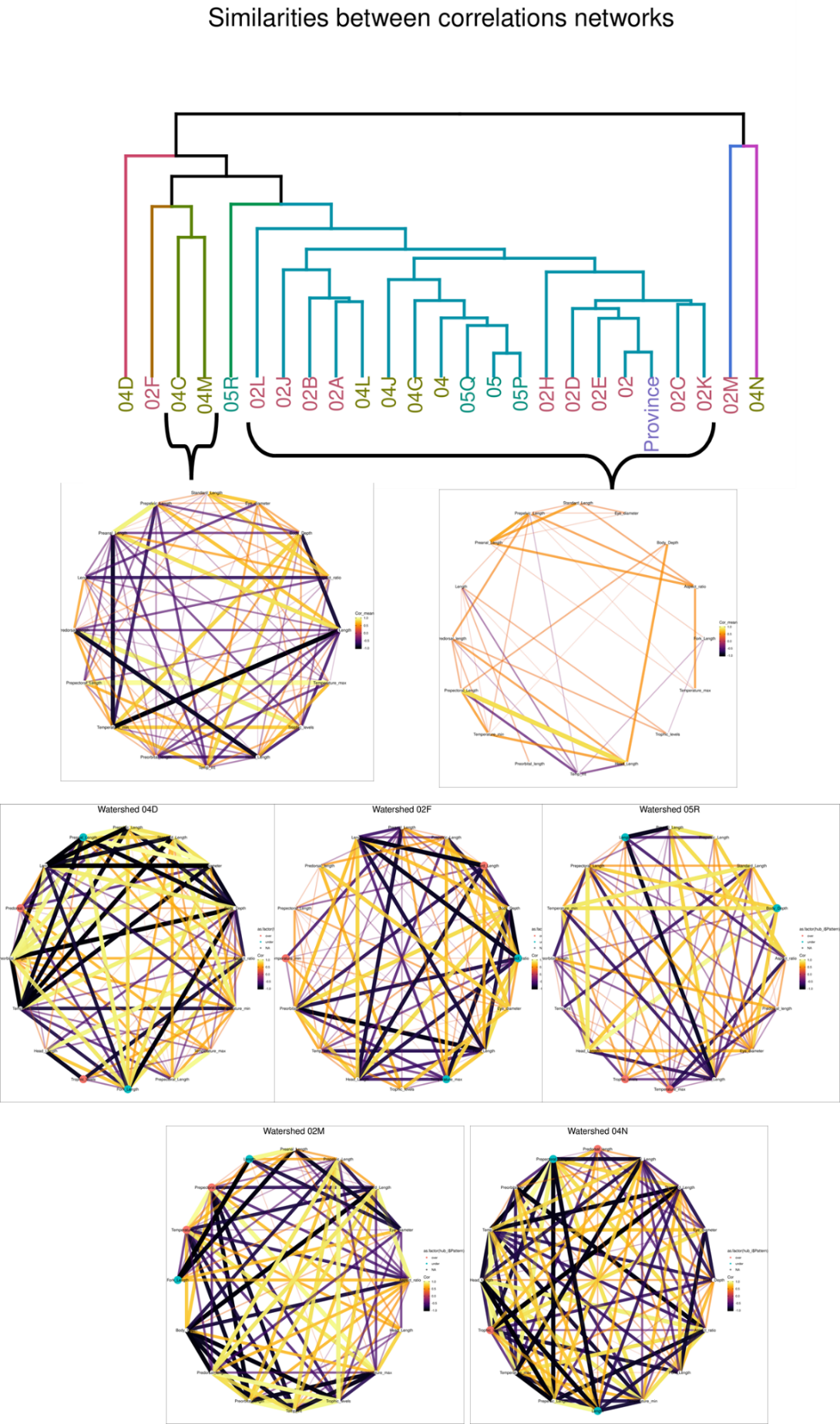

**REFERENCES**

Froese R, Pauly D. 2019. FishBase,version (12/2019). www.fishbase.org
